# Reactivating and Reorganizing Activity-Silent Working Memory Through Two Distinct Mechanisms After Pinging the Brain

**DOI:** 10.1101/2023.07.16.549254

**Authors:** Can Yang, Xianhui He, Ying Cai

## Abstract

Recent studies have proposed that visual information can be maintained in an activity-silent state during working memory (WM), which can be reactivated by task-irrelevant high-contrast visual impulses (i.e., “pinging the brain”). Although pinging the brain has become a popular tool for exploring activity-silent WM in recent years, its underlying mechanisms remain unclear. In the current study, we directly compared the behavioral consequences and neural reactivation effects of context-independent and context-dependent pings to distinguish the noise-reduction and target-interaction hypotheses of pinging the brain. At first, in a behavioral study, we found that, compared with the baseline condition (no ping), only the horizontal context-dependent pings impaired recall performance. In a follow-up electroencephalogram study, our neural decoding results showed that the context-independent pings reactivated activity-silent WM transiently without changing the original WM representations or recall performance. In contrast, the context-dependent pings reactivated activity-silent WM more durably and further reorganized WM information by decreasing the dynamics of items’ neural representations. Notably, only the reactivation strength of the context-dependent pings correlated with recall performance and was modulated by the location of memorized items, with neural representations only being reactivated when both items and pings were presented horizontally. Together, our results provided evidence for two distinct mechanisms underlying pinging the brain, and the ping’s context played a critical role in reactivating and reorganizing activity-silent WM.

## Introduction

Working memory (WM) is the ability to temporarily maintain and manipulate information.^1^ Accumulating studies have demonstrated that humans prioritized WM information processing depending on ongoing tasks.^2,3^ Earlier studies revealed that the neural representations of prioritized memory items (PMIs) were sustained during maintenance, while those of unprioritized memory items (UMIs) dropped to baseline but rebounded as needed.^4,5,6^

Where and how the items are maintained when they are undecodable is always an important theoretical question. Recent studies have explored these processes by pinging the brain with task-irrelevant visual impulses (a.k.a. “pings”). In a pioneering study,^7^ Wolff et al. found that the neural representation of a single item in WM returned to baseline during maintenance. More importantly, they suggested that this item became activity-silent rather than disappearing as a high-contrast white patch could reactivate the item’s neural representation. Subsequently, they extended this finding to multi-item WM, revealing that both PMIs and UMIs could be maintained in the activity-silent state and be reactivated by pinging the brain.^8^ Meanwhile, some studies using computational models also supported the existence of activity-silent WM, suggesting that it could be achieved via transient synapses’ changes.^9,10^ However, the reactivation of WM representations has been challenged by some other studies. For example, a recent study found that pings with ordinal information could only reactivate the sequential information but not the item-specific information.^11^ Another study suggested that pings preferentially reactivated latter items in a sequence but not the former ones.^12^ Currently, pinging the brain has become an important tool for investigating activity-silent WM, and the unclear underlying mechanisms become an increasingly problematic issue.

Pinging the brain was initially introduced as a metaphor for “sonar”, capable of reaching activity-silent memory stored within the brain network and generating a detectable echo of the memorized items.^7^ In this case, the reactivation of a ping could be expected to be related to two main factors. One factor is whether pings share a common context with the memory items, such as overlapped locations. Supporting the crucial role of context in maintaining WM, recent studies have demonstrated that context information (e.g. order or location) is automatically encoded and maintained with content information. ^12,13^ Accordingly, we predict that pings with shared context with memorized items would be more efficient in reactivating activity-silent WM (“context-dependent pings”, like a sonar equipped with an orientation system) and would have the chance to interact with the memorized items (“target-interaction hypothesis”, Figure 1A). In contrast, pings without shared context may work in a more “scattershot” manner (“context-independent pings”). Supporting this view, a recent study reported that pings reduced the across-trial variability of electroencephalograph (EEG) signals, suggesting that pings reactivate activity-silent representations by indirectly reducing the overall neural noise among brain network (“noise-reduction hypothesis”; Figure 1B).^14^ This view has been supported by some other studies which have found that such variability decreasing could also be observed after other types of visual stimuli presentation.^15,16^ The key to distinguishing these two hypotheses is understanding how the context of pings affects reactivations. Unfortunately, most previous studies presented pings totally or partially overlapping with items’ context,^8,17,18^ leaving the answer to this question still open.

**Figure1.**
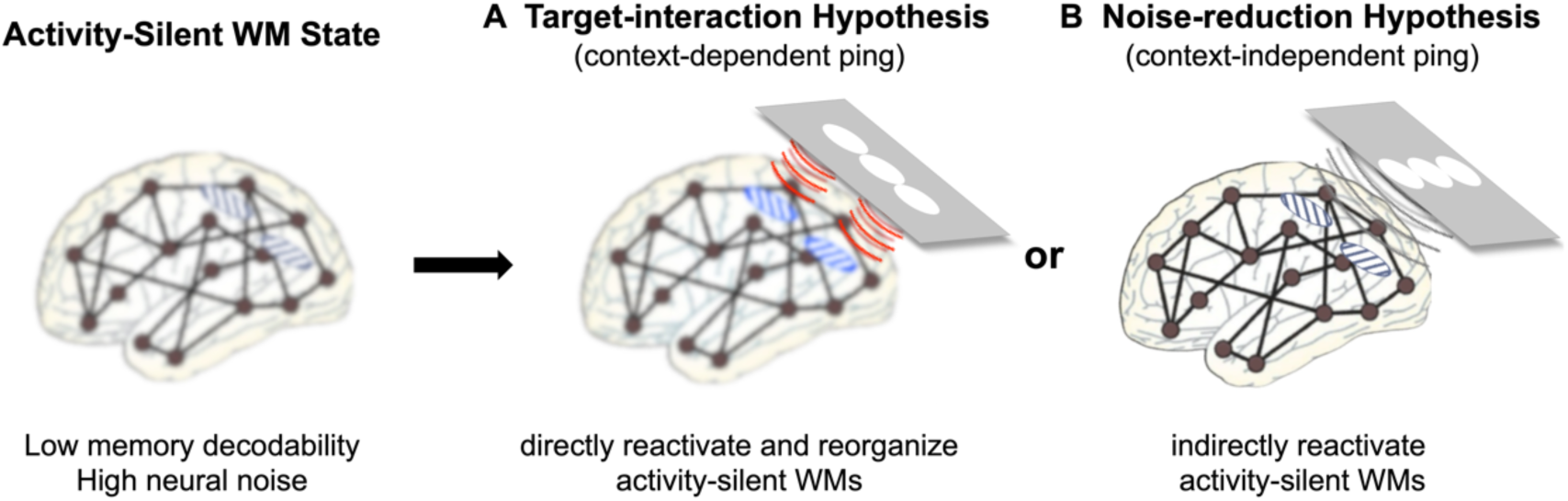
Two hypotheses about the underlying mechanisms of pinging the brain. In the activity-silent WM state, two items were stored within a high-noise brain network in a low decodability condition (left). (*A*) Target-interaction Hypothesis: context-dependent ping directly reactivated activity-silent WM through the shared context (e.g. locations) with the memorized items, with the potential to change the representations of activity-silent WM. (*B*) Noise-reduction hypothesis: context-independent ping indirectly reactivated activity-silent WM by reducing the overall neural noise without interacting with the memorized items.

Another factor is where the memorized items are presented. Previous studies have shown that behavioral performance was better when items were presented along the horizontal meridian than the vertical meridian, such as in letter identification,^19^ spatial and contrast discrimination.^21,22^ This phenomenon, known as horizontal-vertical anisotropy, has also been observed in recent visuospatial WM studies.^23,24^ Researchers usually interpreted it as the spatial representations in the internal priority map were limited by the spatial coding in the visual and oculomotor systems: the spatial coding was more precise along the horizontal axis than that along the vertical axis. Therefore, we predict that context-dependent pings would be even efficient in reactivating neural representations when the memorized items were presented horizontally than vertically. On the contrary, the reactivation effects of context-independent pings would not be modulated by the location of the memorized items. However, there is no direct evidence to test these predictions yet.

In the current study, to directly examine the underlying mechanisms of pinging the brain, we compared neural reactivations in four types of pings: context-dependent pings presented horizontally or vertically and context-independent pings presented horizontally or vertically. Particularly, we tested three critical questions to distinguish the target-interaction hypothesis and the noise-reduction hypothesis. First, whether context-dependent pings are more efficient in reactivating activity-silent WM than context-independent pings. Second, whether only the reactivation effect of context-dependent pings is modulated by their location, specifically if context-dependent, horizontal pings are more efficient than vertical ones. Third, whether only context-dependent pings reorganize the neural representations of activity-silent WM or change the behavioral consequence.

## Results

### Context-dependent, horizontal pings impaired WM performance

One hundred and sixteen participants completed an orientation delayed estimation task adapted from previous studies^4,5,6^ (Figure 2A). The recall order was determined by two retro-cues, with the second one having a 50% probability of switching. Memorized items were presented either horizontally or vertically, and for each participant, three kinds of pings (context-dependent ping/context-independent ping/no ping) were presented before the first recall. When memorized items were presented horizontally, for the response errors in the first recall, the one-way repeated-measures ANOVA revealed a main effect of ping conditions (*F* _(2,108)_ = 3.981, *p* = 0.021, BF_10_ = 1.698). Post-hoc tests demonstrated that the recall error in the context-dependent ping condition was larger than that in the context-independent ping (*t* _(54)_ = 2.708, *p* = 0.024, BF_10_ = 2.883) and no-ping baseline conditions (*t* _(54)_ = 2.041, *p* = 0.087, BF_10_ = 1.178), but there was no difference between the context-independent ping condition and the baseline condition (*t* _(54)_ = 0.667, *p* = 0.506, BF_10_ = 0.183, Figure 2B). In the second recall, the two-way repeated-measures ANOVA results revealed a significant interaction effect between item type and ping condition (*F* _(2,108)_ = 6.636, *p* = 0.002, BF_10_ = 17.754). Further one-way repeated-measures ANOVA results showed a main effect of ping condition for PMI recall (*F* _(2,108)_ = 6.391, *p* = 0.002, BF_10_ = 12.135) but not for UMI recall (*F* _(2,108)_ = 0.675, *p* = 0.511, BF_10_ = 0.108). Post hoc tests showed that the response error for PMI in the context-dependent ping condition was larger than that in both the context-independent ping and baseline conditions (*ts* > 2.415, *ps* < 0.035, BFs > 2.429), but there was no difference between the latter two (*t* _(54)_ = 1.075, *p* = 0.285, BF_10_ = 0.284). These results suggested that the context-dependent, horizontal ping specifically impaired the recall performance of the prioritized items. In contrast, when memorized items were vertical, there were no such error differences among ping conditions in both PMI and UMI recalls (*Fs* < 0.685, *ps* > 0.507, BFs < 0.118, Figure 2B).

**Figure2.**
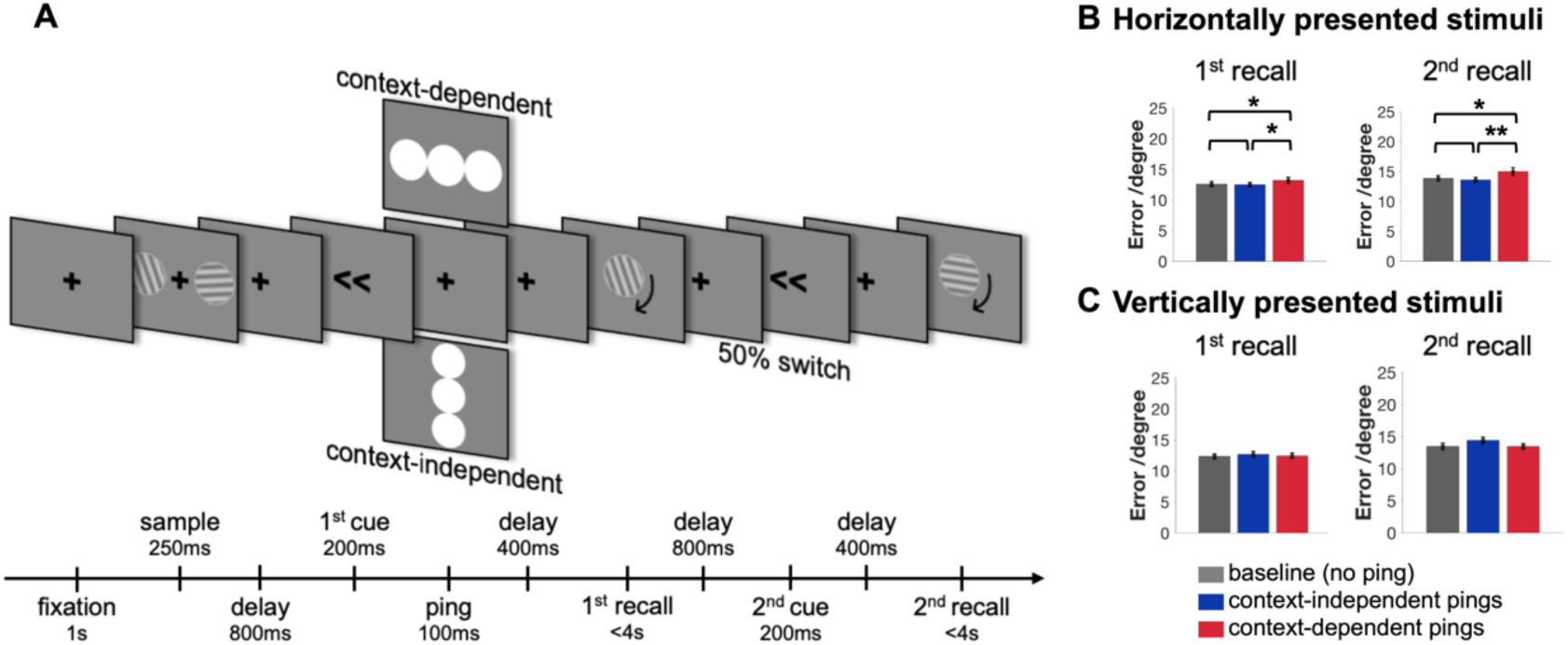
Behavioral Experiment and Results. (*A*) Experiment procedure of the delayed estimation task. The task with horizontally presented items was shown for example. (*B*) The recall error in the two recalls for the three ping conditions when the two orientations were presented horizontally. (*C*) The recall error in the two recalls for the three ping conditions when the two orientations were presented vertically. Error bars indicated the standard error of the mean (SEM). The significant results are noted with asterisks (**: 0.001 < *p* <= 0.01, *: 0.01 < *p* <= 0.05).

### Both context-dependent pings and context-independent pings reactivated prioritized memory items

Another Sixty-three participants completed a similar orientation delayed estimation task while recording the EEG signals, except that the experimental procedures were slightly changed to increase the neural decoding efficiency (refer to a previous study,^8^ see more details in the methods, Figure 3A). In all ping conditions, both PMIs and UMIs could be decoded during the sample and early delay periods, while their decoding strengths decayed to baseline during the late delay period, suggesting that both PMIs and UMIs were maintained in an activity-silent state. Then, the neural representation of PMIs rebounded after the context-dependent, horizontal pings and the two context-independent pings, but not after the context-dependent, vertical pings. In contrast, UMIs did not exhibit any reactivation after any ping condition (Figure 3B, see more details about neural decoding and cluster-based statistics in the methods).

**Figure3.**
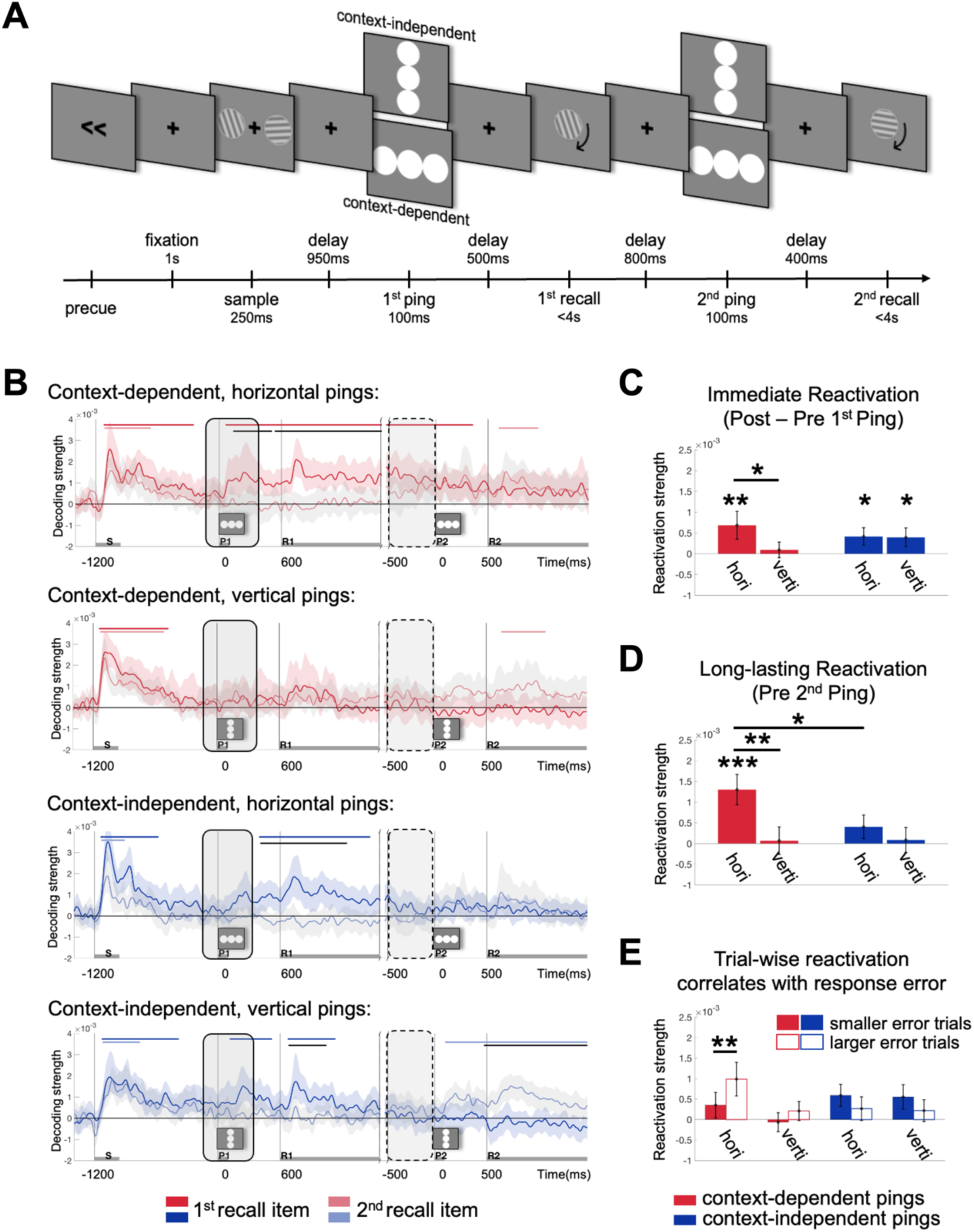
Task procedure in EEG experiment and neural decoding and reactivation results. (*A*) Experiment procedure of the delayed estimation task. Gratings could be presented along the horizontal or vertical axis (only the horizontal case was shown here). (*B*) Decoding strengths of the first recall item (PMI) and the second recall item (UMI) under the four ping conditions. The blue and red bars on the top of each plot indicated clusters with significant decoding strengths and the black bars indicated clusters with significant decoding differences between the two items (*p* < 0.05). Error shading along the decoding strength line indicated a 95% confidence interval. Gray bars on the X-axis indicated the duration of the sample (S), the first ping (P1), the first recall (R1), the second ping (P2), and the second recall (R2). Transparent gray areas with solid borders indicated the time window used to estimate the immediate reactivation strengths, and transparent gray areas with dashed borders indicated the time window used to estimate the long-lasting reactivation strengths. (*C*) Immediate reactivation strengths of PMI under the four ping conditions. (*D*) Long-lasting reactivation strengths of PMI under the four ping conditions. (*E*) Immediate reactivation strengths of PMI in the half trials with smaller recall errors and larger recall errors under the four ping conditions. Error bars indicated the standard error of the mean (SEM). The significant results are noted with asterisks (***: *p* < 0.001, **: 0.001 < *p* < = 0.01, *: 0.01 < *p* <= 0.05).

To estimate the immediate reactivation strength, we subtracted the pre-ping decoding strengths (the averaged decoding strengths from −100 ∼ 0 ms before the ping) from the post-ping decoding strengths (the averaged decoding strengths from 100 ∼ 400 ms after the ping).^25^ Our analyses focused on the first ping period when both the PMI and UMI were task-relevant and simultaneously maintained, and the results showed that only the reactivation effect of context-dependent pings was modulated by the location of memorized items. That is, PMI was only reactivated after the context-dependent, horizontal pings but not the context-dependent, vertical pings (context-dependent, horizontal ping: *p* = 0.011, context-dependent, vertical ping: *p* = 0.305, difference: *p* = 0.046). In contrast, context-independent pings evoked comparable immediate reactivations regardless of the locations per se (context-independent, horizontal ping: *p* = 0.021, context-independent, vertical ping: *p* = 0.032, difference: *p* = 0.526). However, no differences were found between context-dependent and context-independent pings when they were presented either horizontally or vertically (*ps* > 0.143; Figure 3C; all *p* values were obtained by bootstrapping procedures, see more details in the methods).

Besides, to quantify the long-lasting reactivation effect, we averaged the decoding strengths from −500 to 0 ms before the second ping (i.e., the interval after the first recall and before the second ping). As we expected, our results showed that only the context-dependent, horizontal pings evoked a long-lasting reactivation effect (*p* < 0.001, other three ping conditions: *ps* > 0.073, differences: *ps* < 0.025, Figure 3D), suggesting that both the shared context evoked a more sustained neural response and it was modulated by the locations of memorized items per se.

We also tested the reactivations using voltage signals among anterior, middle brain areas, and alpha power patterns among posterior areas, but found no reliable reactivations (see more details in the Supplemental Information, Figure S1 and S2). Then, all the following reactivation analyses only focused on voltage signals among posterior electrodes.

### Larger reactivations after the context-dependent pings related to larger recall errors

For the significant reactivation effects, we further tested the relationship between reactivation strengths and behavioral performance in different ping conditions. We compared the immediate reactivation strengths between the two subgroups of trials (a.k.a. larger errors vs. smaller error trials, based on the median of recall errors). Consistent with the specific long-lasting reactivations, only in the context-dependent, horizontal ping condition, we found that the neural reactivation was stronger in trials with larger recall errors compared to those with smaller recall errors (*p* = 0.005, other ping conditions, *ps* > 0.134, Figure 3E).

### Context-dependent pings stabilized the neural representations of the prioritized memory items

For the significant reactivation effects, we also conducted cross-time decoding analyses to examine whether and how different types of pings affected the dynamics of activity-silent WM storage. Our results revealed that neural representations of PMIs were highly dynamic during the sample and maintenance periods, which were comparable across all four ping conditions. However, the cross-time neural representation strengths were significant only after the context-dependent, horizontal ping (cluster-corrected, *p* < 0.05; Figure 4A). Consistently, the generalization index, which was calculated as the averaged cross-time decoding strengths wherein the decoder was trained after the ping and tested on the following recall period, was only significant after the context-dependent, horizontal ping (*p* = 0.003) whereas there were no such effects in other ping conditions (*ps* > 0.101, see more details in the methods). Moreover, this difference was significant when context-dependent pings were presented horizontally compared to vertically (*p* = 0.042; Figure 4B). These results suggested that only the context-dependent pings reorganized the neural representations of PMIs, which decreased the dynamics of the neural representations.

**Figure4.**
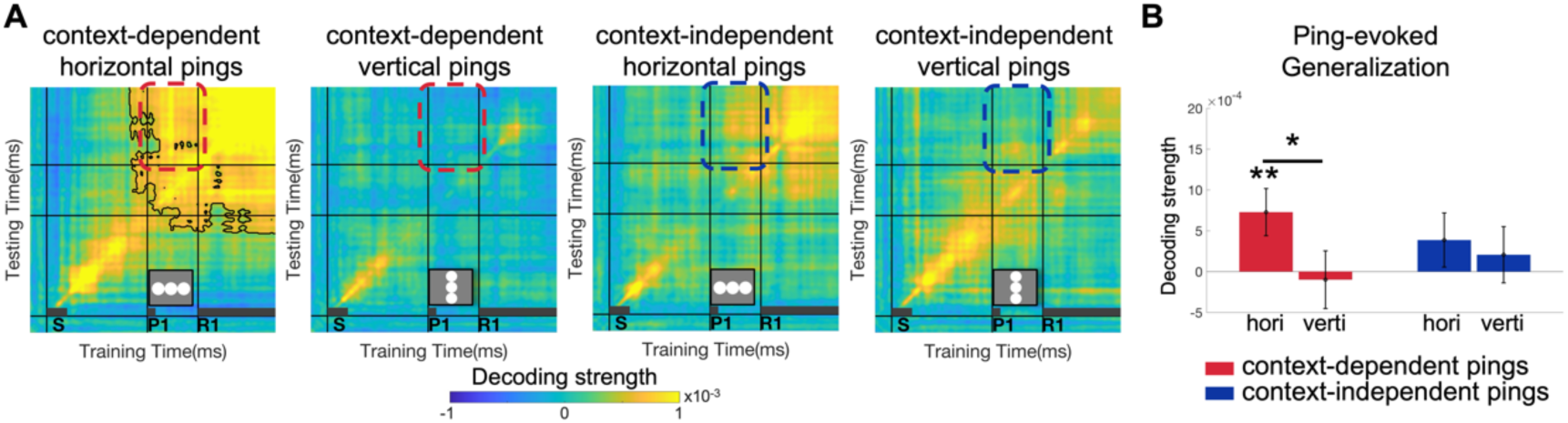
Cross-time decoding and representational generalization results in different ping conditions. (*A*) Cross-time decoding strengths of the first recall item (PMI) under the four ping conditions. The black lines in the decoding matrix outlined clusters with significant decoding strength (*p* < 0.05). Gray bars on the X-axis indicated the duration of the sample (S), the first ping (P1), and the first recall (R1). Dashed boxes in red and blue indicated the time window used to estimate the generalization strengths. (*B*) Ping-evoked generalization strengths of the first recall item under the four ping conditions. Error bars indicated the standard error of the mean (SEM). The significant results are noted with asterisks (**: 0.001 < *p* <= 0.01, *: 0.01 < *p* <= 0.05).

### Noise reduction was not a ping-specific consequence

For the significant reactivation effects, we directly tested the noise-reduction hypothesis by estimating changes in across-trial variability, which was used to index the level of neural noise.^14^ The noise reduction index was estimated in four ping conditions after each visual stimulus was presented (i.e. the presentation of fixations, sample arrays, pings, and probe displays). However, our results revealed that neural noise generally reduced after all the visual stimuli and did not correlate with reactivation strength in any ping condition (see more details in the Supplemental Information, Figure S3).

## Discussion

Our study suggested that context-dependent and context-independent pings reactivated activity-silent WMs through two distinct mechanisms. The context-independent ping reactivated the neural representations without changing the original WM representations or impacting recall performance, and its reactivation was similar regardless of the location of memorized items per se. In contrast, the context-dependent ping reactivated activity-silent WMs more durably and also reduced the neural representations’ dynamics, which correlated with impaired recall performance. Notably, the reactivation effect of the context-dependent ping was modulated by the location of memorized items.

Supporting the noise-reduction hypothesis, our results revealed that context-independent pings that did not overlap with the locations of memorized items could reactivate the activity-silent WM. Compared to previous studies using pings that either totally or partially overlapped with memorized items’ locations,^8,18^ our results clearly demonstrated that the ping itself can reactivate activity-silent WM without relying on any shared contextual information with items. Consistently, although our across-trial variability results revealed the noise-reduction after ping was not ping-specific, it could be a reasonable mechanism to explain the reactivation effects as previous studies suggested^14^. Moreover, our cross-time decoding results found that context-independent ping neither altered the dynamics of activity-silent WM representation nor affected the recall performance. These results supported the original “sonar” metaphor of pinging the brain, which proposed pings as a “detection” tool and would not cause any changes of the targets per se.

Meanwhile, our results also provided evidence for the target-interaction hypothesis of pinging the brain in the context-dependent ping conditions. Firstly, we found that compared with context-independent pings, context-dependent pings reactivated WM representations more durably and its reactivation effect was modulated by location of memorized items. On one hand, these results highlighted a more efficient pathway of pinging the brain through shared contextual information. These results were supported by some recent studies that also emphasized the importance of the contextual information of pings.^13,26^ For example, recent studies indicated the neural representations of items’ context and content were integrated during maintenance and only the strength of context-dependent neural representation predicted recall performance. Accordingly, another latest study revealed that, during a visual search task, pings at locations with a higher probability of being tested reactivated the neural representation (“prioritized location”).^27^ On the other hand, the location effects of context-dependent pings were consistent with recent studies reporting the horizontal-vertical anisotropy during VWM,^23,24^ which have revealed that the items presented along the horizontal meridian obtained better behavioral performance than those along the vertical meridian. Both sets of evidence suggested that the overlapped context made the pings work more efficiently via the internal priority map.

Secondly, we observed that context-dependent pings reorganized the neural representations by reducing their dynamics, which impaired behavioral performance. Specifically, compared with the dynamic representations after context-independent pings, there were shared neural representations after context-dependent pings and during recall period, indicating that context-dependent pings facilitated the neural representation transformation from WM maintenance to response preparation. Consistently, recent studies have shown that individuals would proactively generate item-specific action plans and integrate them with WM content during the late delay period, leading to faster behavioral responses.^28^ Besides, our results were consistent with previous findings that the highly dynamic changes in neural representations have a positive effect on recall performance. For instance, some recent intracranial EEG studies employing deep learning and pattern similarity analyses have demonstrated that such dynamic neural representations during WM reflected the transformation from maintaining visual details to abstract information, benefiting both WM and long-term memory.^29,30^ However, some other studies found that trials with stronger neural representations after ping were accompanied by higher recognition accuracy^8^ or reported no behavioral relevance of item-specific reactivations.^11^ We suspected these inconsistent behavioral consequences may be attributed to different experimental settings. In our main behavioral experiment, which minimized confounding factors such as expectation and adaptation effects by using retro-cues and an interleaved design, we observed overall worse recall performance specifically after context-dependent, horizontal pings. However, in our EEG experiment, we used pre-sample cues and block design to maximize SNR for neural decoding like most previous studies. Although we did not observe behavioral changes between ping conditions, we observed larger recall errors after higher reactivations in context-dependent horizontally presented ping. In this case, the relationship between reactivation effects and behavioral performance was consistent while in different ways. Nevertheless, this issue should be examined further in future studies.

Furthermore, our results clarified several conflicts in the understanding of the activity-silent WM storage. One is that our results localized the activity-silent WM representations in the relevant sensory cortices. Our results were consistent with previous studies reporting neural reactivations in the posterior electrodes^8,18^ and helped to reconcile conflicting results that signals were obtained and examined among different electrodes.^31,32^ This also suggested that neural coding results using whole-brain signals should be interpreted with caution.^11,33^ The uniqueness of reactivation effects in the posterior sensory cortex aligned with some previous studies. For instance, the sensory-recruitment theory of VWM posited the critical role of the sensory cortices in maintaining item-specific WM information,^34,35,36^ and recent studies have proposed that these representations were susceptible to external visual input during maintenance.^37^

Another noteworthy finding in our study is that neither context-dependent nor context-independent pings reactivated the unprioritized items, implying that while both prioritized and unprioritized items were undecodable in the activity-silent state, task priority still modulated the WM storage states. These results were consistent with previous studies that have reported weaker reactivations for unprioritized items^8^ and that the unprioritized items are more resistant to external interruptions during maintenance.^38^ In addition, a recent study suggested that WM items are reorganized into different activity-silent states depending on their sequence context and can be reactivated at different latencies during retention after pings.^12^ However, some other studies found that unprioritized items can also be reactivated by pinging the brain ^8,12^ or by transcranial magnetic stimulation (TMS).^39^ We propose that these inconsistencies are due to different settings of pings across studies. For example, pings can be high-contrast or pure white impulses,^7,8^ be a single large visual stimulus covering almost the whole screen^11,12^ or consist of several small visual patches wholly or partially covering the memorized items’ locations.^18,27^ Besides, the duration of pings can also differ from 100 ms to 200 ms.^8,27^ Thus, the efficiency of pings could be a salient confounding factor and should be considered in future studies. Moreover, the discrepancy between our findings and those of the TMS study may indicate that pinging and TMS operate through distinct mechanisms, and other investigations can explore these differences in future studies.

Although our study has provided valuable insights into the underlying mechanisms of pinging the brain, several limitations should be noted. First, although our results suggested that context-dependent and context-independent pings evoked reactivations through distinct pathways, future studies are needed to determine whether these two mechanisms act independently or whether the reactivation effect of context-dependent pings is a combined result of both mechanisms. Second, our EEG results provided evidence for reactivations among the posterior electrodes while sustaining neural representations in frontal areas; future studies can use imaging techniques with both high temporal and high spatial resolution (such as magnetoencephalography) to further explore how pings affect the dynamics of neural representations across the whole brain. Additionally, to directly verify the neural dynamic changes after pings, it would be helpful to include a condition without pings and conduct cross-condition decoding in future studies.

In summary, the current study directly examined the context and location effects of pinging the brain by comparing their neural reactivation effects and behavioral changes. Our results clarified two distinct underlying mechanisms of pinging the brain: the context-independent ping reduces the sensory-network representation noise and evokes a transient neural echo without changing the original WM maintenance or behavioral performance, whereas the context-dependent ping accesses and interacts with activity-silent WM through an internal priority map that is organized by contextual information and items’ location, resulting in impaired WM performance. These results offer valuable insights for future studies employing pings as a tool to investigate activity-silent WM and enhance our understanding of the activity-silent WM process.

## Methods

### Experimental model and subject details

This study was approved by the Institutional Review Board of the Department of Psychology and Behavioral Sciences, Zhejiang University, and complies with all relevant ethical regulations. All participants gave written informed consent and were compensated for their participation (30 CNY/h). Participants had normal or corrected-to-normal vision and no reported neurologic or psychiatric disease history.

### Participants in behavioral experiment

Sixty right-handed volunteers (33 females; M_age_ = 21.17 years, SD_age_ = 2.49 years) participated in the behavioral experiment using horizontally-presented items, and fifty-six right-hand volunteers (33 females; M_age_ = 22.80 years, SD_age_ = 3.05 years) participated in the same behavioral experiment while using vertically-presented items. The sample size for the horizontal-item task was estimated using G*power 3.1^40^ based on a pilot experiment (with α set to 0.05; and power set to 0.8), and the sample size for the vertical-item task was estimated based on the horizontal experiment using a similar method. Five participants were excluded from the group experimenting with horizontally-presented items and six participants were excluded from the group experimenting with vertically-presented items due to poor behavioral performance (RT or recall error was more than 3 SDs away from the group means).

### Participants in EEG experiment

Another sixty-three right-handed volunteers (25 females, M_age_ = 21.89 years, SD_age_ = 2.68 years) participated in the EEG experiment. The sample size was determined based on previous EEG studies.^8,17^ Six participants were excluded from all subsequent EEG analyses due to poor signal quality (3 subjects, more than 30% bad trials) or poor behavioral performance (3 subjects, reaction time or recall error was more than 3 SDs away from the group means). In the EEG analyses, all decoding analyses focused on the first 1 s after each response probe since the mean reaction time were approximately 1 s for both recalls (recall 1: mean ± SD = 1.06 ± 0.33 s; recall 2: mean ± SD = 1.15 ± 0.36 s).

### Task procedure in behavioral experiment

Participants completed an orientation delayed estimation task in the behavioral experiment (Figure 2A). At the beginning of each trial, a 1000 ms fixation was presented. Then, two oriented gratings appeared for 250 ms, either to the left and right (horizontally presented) or above and below (vertically presented) the centered fixation (6° eccentricity). These gratings were sine-wave patterns with a contrast of 20%, a diameter of 6°, and a spatial frequency of 0.65 cycles per degree. The orientations were selected randomly from a pool of 36 options, ranging from 0° to 180° in 5° increments, and with a minimum difference of 20° between any two orientations in the same trial. After a delay of 800 ms, the first retro-cue was presented for 200 ms, indicating the item to be recalled first. Following the first retro-cue, the ping was presented for 100ms. In the context-dependent ping conditions, three white circle impulses (6° in diameter) were presented, with two on the sides overlapping with the locations of the memorized items during the sample period; in the context-independent ping conditions, pings involved the same three impulses but they were rotated by 90° and did not overlap with the memorized items; in the baseline condition without any ping, a fixation dot was presented at the center of the screen. The centered visual impulse was included to emphasize the task irrelevance of the pings. After another 400 ms delay, participants were required to recall the prioritized item by moving the mouse to select the corresponding orientation value, with a maximum response window of 4 seconds. The second retro-cue appeared 800 ms after the first response, with a 50% probability of switching. In the switch condition, the item to be recalled second was the UMI in the first recall; in the stay condition, the item to be recalled second was the same as the first recalled item (a.k.a. PMI). After an additional 400 ms delay, participants were tasked to recall the second-cued item in the same manner. For half of the participants, two orientations were always presented horizontally, and each ping condition was divided into two blocks. The order of the blocks was counterbalanced across participants. The other half of the participants completed trials with orientations presented vertically, with the same ping condition settings. Thus, each participant completed all six blocks within approximately 60 minutes, consisting of 104 trials for each ping condition.

### Task procedure in EEG experiment

Participants completed a similar delayed estimation task as in the behavioral experiment, except that the recall order for two orientations was instructed at the beginning of each trial and two pings were presented during the maintenance before each recall (Figure 3A). For half of the participants, two orientations were always presented horizontally, and context-dependent and context-independent pings were organized in two separate sessions 48 hours apart. The recall order of two orientations remained consistent within each session and was counterbalanced across participants. The order of context-dependent/independent ping conditions were also counterbalanced across participants. The other half of the participants completed trials with orientations presented vertically, with the same ping condition settings. Thus, in each session, each participant completed all the trials from one ping condition within approximately 90 minutes, consisted of 540 trials across 9 blocks. Such design for four ping conditions aimed to maximize the signal-to-noise ratio (SNR) for neural representation analyses.^8^ All the experiments were programmed and executed using MATLAB (MathWorks) and the Psychophysics Toolbox extension (http://psychtoolbox.org). Visual stimuli were displayed on a 17-inch CRT monitor with a refresh rate of 100 Hz and a resolution of 1024 x 768 pixels. Participants were seated in a dark room and maintained a viewing distance of approximately 60 cm from the screen. The background color of the screen was set to gray (RGB = 128, 128, 128) throughout the experiment to minimize potential visual distractions.

### EEG acquisition and preprocessing

During the EEG experiment, neural signals was recorded using a Biosemi Active Two recording system with a sampling rate of 1024 Hz. The 64 Ag/AgCl electrodes were positioned according to the extended 10–20 system, with a reference of a common mode sense electrode (CMS/DRL). During the recording, the impedances of all the channels were kept below 15 kΩ. EEG data preprocessing and analysis were conducted using the functions in the EEGLAB toolbox^41^ and customized MATLAB scripts. Horizontal eye movements were monitored by the horizontal electrooculogram (EOG) placed at the outer canthus of the two eyes, and vertical eye movements were monitored by the vertical EOG placed above and below the left eye. Raw EEG data were first downsampled to 256 Hz, bandpass filtered (0.1 ∼ 40 Hz), and segmented into two epochs for each trial (from −2 s to 2 s relative to the onset of the first and second pings; we segmented the two epochs separately since the recalls were self-paced). After segmentation, bad EEG channels were identified by visual inspection and then interpolated using the “spherical” method in EEGLAB. Baseline removal was conducted by subtracting the averaged activity during the 200 ms prestimulus interval. Eye blinks and muscle artifacts were identified through independent component analysis (ICA) and removed from the data. Finally, the post-ICA data were carefully visually inspected for any potential remaining artifacts, which were then rejected. As a result, 10.87% (SD = 6.31%) of the trials were excluded, and approximately 480 trials were kept for each ping condition (context-dependent, horizontal ping: Mean ± SD = 473.57 ± 29.45; context-dependent, vertical ping: Mean ± SD = 498.22 ± 18.46; context-independent, horizontal ping: Mean ± SD = 495.85 ± 22.56; context-independent, vertical ping: Mean ± SD = 460.77 ± 43.94).

### Data analysis for behavioral performance

In the behavioral experiment, for the responses during the first recall, we conducted one-way repeated-measures ANOVAs to examine whether the three ping conditions led to different behavioral consequences. If the main effect was significant, we conducted further post hoc tests to check the differences between the three ping conditions. For the responses during the second recall, we first conducted a two-way repeated-measures ANOVA, with the within-subject factors of ping condition (context-dependent ping/context-independent ping/baseline) and item type (PMI/UMI). If the interaction effect was significant, we further conducted one-way repeated-measures ANOVAs and post hoc tests for PMI and UMI recall separately to check the differences between the three ping conditions. All reported *p* values were Holm‒Bonferroni multi-comparison corrected. We did these analyses for the horizontal and vertical versions separately. In the EEG experiment, similar ANOVA and post-hoc t-tests were conducted. However, there was no difference in recall errors between any ping condition (*ps* > 0.864, BF_10s_ < 0.514).

### Estimation of the decoding strength of neural representations

We estimated the neural representations of PMIs and UMIs in each ping condition separately referring to previous studies.^8,12^ For each time point, we extracted the voltage pattern across 17 parietal and occipital channels (P7, P5, P3, P1, Pz, P2, P4, P6, P8, PO7, PO3, POz, PO4, PO8, O1, Oz and O2), and normalized these signals by subtracting their average value. Then, we aggregated the voltage patterns at 5 consecutive time points (including the time point of interest and its adjacent four time points) for neural decoding analyses. Refer to previous studies, including both spatial and temporal information can increase the sensitivity of neural decoding.^11,17^ Next, we used the Mahalanobis distance^42^ method and the leave-one-trial-out cross-validation approach to estimate the posterior decoding strength of neural representations. For each trial, the data from that trial served as the testing data, and the data from all the remaining trials served as the training dataset. The training trials were divided into 12 bins based on their orientations relative to the test trial. The neural activity patterns of the trials labeling the same bin were averaged, and the Mahalanobis distances were calculated between the spatiotemporal neural activity pattern of the test trial and of each training bin. To estimate the decoding strength, 12 sign-reversed Mahalanobis distances were multiplied by the cosine of the center angle of each bin. This procedure was repeated for all trials and all time points, and the decoding strengths were averaged over trials, smoothed over time with a Gaussian smoothing kernel (SD = 16 ms) and then averaged across participants. Finally, we utilized a nonparametric sign-permutation test to identify the time period in which the decoding strengths were significant.^43^ For each time point in each epoch, the sign of the decoding strength was randomly flipped for every participant, and the average was calculated. After 5,000 permutations, we obtained a null distribution of decoding strengths and the threshold for significant decoding (with α set to 0.05, two-tailed). If two consecutive time points exceeded the threshold for significant decoding, we marked them as cluster candidates and aggregated their decoding strengths. Taking the decoding strengths of the 5,000 permutations for each time point together, we also obtained two null cluster-based distributions for the smallest and highest decoding strengths. *P* values referred to the proportion of the real cluster-based decoding strengths larger than the maximum null distribution or smaller than the minimum null distribution. Some recent studies suggested that item-specific information could be sustainably decoded from posterior alpha power activities during WM maintenance and were not activity-silent.^14,44^ Thus, we also conducted a similar decoding analysis based on posterior alpha power activities (8 ∼ 12 Hz). Meanwhile, to further explore whether the storage and reactivation of activity-silent WM is location-specific on brain regions, we also conducted similar decoding analyses among 17 anterior (FP1, FPz, FP2, AF7, AF3, AFz, AF4, AF8, F7, F5, F3, F1, Fz, F2, F4, F6 and F8) and mid-central (FC3, FC1, FCz, FC2, FC4, C5, C3, C1, Cz, C2, C4, C6, CP3, CP1, CPz, CP2 and CP4) electrodes. However, no reactivation effects were observed using alpha power signals (Figure S1) or using voltage singles from other areas (Figure S2). Therefore, all the following reactivation analyses focused on voltage signals among posterior electrodes.

### Estimation of the reactivation strength after pings

We used the bootstrapping procedure to examine whether the reactivation strengths for the PMIs and UMIs were significant in each ping condition. Specifically, we randomly resampled all the participants with replacement 5,000 times to obtain a null distribution of reactivation strengths for each ping condition, and the *p* values referred to the proportion of the null distribution less than zero. Moreover, we further examined the context and location effects of pings on the reactivation strengths. To examine the context effect of pings, we used a similar bootstrapping procedure to compare the reactivation strength difference between context-dependent and context-independent pings when they were presented in the same horizontal or vertical way. Specifically, we subtracted the null-distributions of reactivation strengths for context-dependent and context-independent ping conditions to create a null-distribution of difference, and the *p* value referred to the proportion of the null-distribution of difference that was greater than or less than zero. To examine the location effect of pings, we compared the reactivation strengths between horizontal and vertical pings when they were in the same context-dependent or context-independent conditions using the similar bootstrapping procedures.

### Estimation of the dynamic changes in neural representations after pings

For the significant reactivation effects, we also conducted cross-time decoding analyses to examine whether and how different types of pings affected the dynamics of activity-silent WM storage. We used a similar approach to estimate the decoding accuracy, except that we calculated the Mahalanobis distances between the spatiotemporal neural activity patterns across different time points. To assess the significance of the cross-time decoding strengths, we performed the same nonparametric sign-permutation test (with 5,000 permutations). *P* values referred to the proportion of the real cluster-based decoding strengths larger than the maximum null distribution cutoff. To further quantify the representational generalization, we averaged the cross-time decoding strengths wherein the decoder was trained after the ping and tested on the following recall period. Then, we used the same bootstrapping procedure to examine whether the generalization strengths were significant in each ping condition. Moreover, to examine the context and location effects of pings, we compared the differences in generalization strengths between context-dependent and -independent ping conditions, as well as between horizontal and vertical pings, using the same bootstrapping procedures.

### Estimation of the neural noise reduction after pings

First, we detrended the raw voltage signal of each electrode for each trial to remove the signal drift, then the variability was computed as the standard deviation of the voltage across all trials at each time point for each electrode. Second, we computed the percentage change of the variability relative to the baseline (−100 ms ∼ 0 ms before the ping and 100 ms ∼ 400 ms after the ping). The noise reduction index was estimated for four ping conditions separately and statistically tested using similar nonparametric sign-permutation test methods. Besides, we did similar analyses after each visual stimulus was presented (i.e. the presentation of sample and probe displays) to test the specificity of after-ping neural activity changes.

## Data and code availability

- Data is deposited on the Open Science Framework at https://osf.io/af54w/
- All original code used for analysis of the data is available on the Open Science Framework at https://osf.io/af54w/
- Any additional information required to reanalyze the data reported in this paper is available from the lead contact upon request.

## Ackonwledgments

This work was supported by the National Natural Science Foundation of China (32100851), the Ministry of Education of Humanities and Social Science Project (21YJC19002), the Zhejiang Provincial Natural Foundation Grant (LQ21C090005), the Fundamental Research Funds for the Central Universities (2021FZZX001-06), and the Research of Basic Discipline for the 2.0 Base of Top-notch Students Training Program, the Ministry of Education of China (20211033).

## Author contributions

Conceptualization, C. Y., and Y. C.; methodology, C. Y., and Y. C.; formal analysis, C. Y., and Y. C.; investigation, C. Y., and Y. C.; writing – original draft, C. Y., and Y. C.; writing – review & editing, C. Y., Y. C., and X. -H. H.; visualization, C. Y., Y. C., and X. -H. H.; funding acquisition, Y. C.

## Supplemental information

Document S1. Figure S1-S3

## References

1. Baddeley, A. (2003). Working memory: Looking back and looking forward. Nature Reviews Neuroscience, 4(10), 829–839. 10.1038/nrn1201

2. Gazzaley, A., & Nobre, A. C. (2012). Top-down modulation: Bridging selective attention and working memory. Trends in Cognitive Sciences, 16(2), 129–135. 10.1016/j.tics.2011.11.014

3. Myers, N. E., Stokes, M. G., & Nobre, A. C. (2017). Prioritizing Information during Working Memory: Beyond Sustained Internal Attention. Trends in Cognitive Sciences, 21(6), 449–461. 10.1016/j.tics.2017.03.010

4. LaRocque, J. J., Lewis-Peacock, J. A., Drysdale, A. T., Oberauer, K., & Postle, B. R. (2013). Decoding Attended Information in Short-term Memory: An EEG Study. Journal of Cognitive Neuroscience, 25(1), 127–142. 10.1162/jocn_a_00305

5. Lewis-Peacock, J. A., Drysdale, A. T., Oberauer, K., & Postle, B. R. (2012). Neural Evidence for a Distinction between Short-term Memory and the Focus of Attention. Journal of Cognitive Neuroscience, 24(1), 61–79. 10.1162/jocn_a_00140

6. Lewis-Peacock, J. A., & Postle, B. R. (2012). Decoding the internal focus of attention. Neuropsychologia, 50(4), 470–478. 10.1016/j.neuropsychologia.2011.11.006

7. Wolff, M. J., Ding, J., Myers, N. E., & Stokes, M. G. (2015). Revealing hidden states in visual working memory using electroencephalography. Frontiers in Systems Neuroscience, 9. 10.3389/fnsys.2015.00123

8. Wolff, M. J., Jochim, J., Akyürek, E. G., & Stokes, M. G. (2017). Dynamic hidden states underlying working-memory-guided behavior. Nature Neuroscience, 20(6), 864–871. 10.1038/nn.4546

9. Barak, O., & Tsodyks, M. (2014). Working models of working memory. Current Opinion in Neurobiology, 25, 20–24. 10.1016/j.conb.2013.10.008

10. Mongillo, G., Barak, O., & Tsodyks, M. (2008). Synaptic Theory of Working Memory. Science, 319(5869), 1543–1546. 10.1126/science.1150769

11. Fan, Y., & Luo, H. (2023). Reactivating ordinal position information from auditory sequence memory in human brains. Cerebral Cortex, 33(10), 5924–5936. 10.1093/cercor/bhac471

12. Huang, Q., Zhang, H., & Luo, H. (2021). Sequence structure organizes items in varied latent states of working memory neural network. eLife, 10, e67589. 10.7554/eLife.67589

13. Yu, Q., Teng, C., & Postle, B. R. (2020). Different states of priority recruit different neural representations in visual working memory. PLOS Biology, 18(6), e3000769. 10.1371/journal.pbio.3000769

14. Barbosa, J., Lozano-Soldevilla, D., & Compte, A. (2021). Pinging the brain with visual impulses reveals electrically active, not activity-silent, working memories. PLOS Biology, 19(10), e3001436. 10.1371/journal.pbio.3001436

15. Arazi, A., Gonen-Yaacovi, G., & Dinstein, I. (2017). The Magnitude of Trial-By-Trial Neural Variability Is Reproducible over Time and across Tasks in Humans. eNeuro, 4(6). 10.1523/ENEURO.0292-17.2017

16. Churchland, et al. (2010). Stimulus onset quenches neural variability: A widespread cortical phenomenon. Nature Neuroscience, 13(3), 369–378. 10.1038/nn.2501

17. Wolff, M. J., Jochim, J., Akyürek, E. G., Buschman, T. J., & Stokes, M. G. (2020). Drifting codes within a stable coding scheme for working memory. PLOS Biology, 18(3), e3000625. 10.1371/journal.pbio.3000625

18. Kandemir, G., Wolff, M. J., Karabay, A., Stokes, M. G., Axmacher, N., & Akyürek, E. G. (2022). Concurrent maintenance of both veridical and transformed working memory representations [Preprint]. Neuroscience. 10.1101/2022.10.28.514218

19. Mackeben, M. (1999). Sustained focal attention and peripheral letter recognition. Spatial Vision, 12(1), 51–72. 10.1163/156856899X00030

20. Carrasco, M., P.Talgar, C., & Cameron, E. L. (2001). Characterizing visual performance fields: Effects of transient covert attention, spatial frequency, eccentricity, task and set size. Spatial Vision, 15(1), 61–75. 10.1163/15685680152692015

21. Baldwin, A. S., Meese, T. S., & Baker, D. H. (2012). The attenuation surface for contrast sensitivity has the form of a witch’s hat within the central visual field. Journal of Vision, 12(11), 23–23. 10.1167/12.11.23

22. Greenwood, J. A., Szinte, M., Sayim, B., & Cavanagh, P. (2017). Variations in crowding, saccadic precision, and spatial localization reveal the shared topology of spatial vision. Proceedings of the National Academy of Sciences, 114(17), E3573– E3582. 10.1073/pnas.1615504114

23. Montaser-Kouhsari, L., & Carrasco, M. (2009). Perceptual asymmetries are preserved in short-term memory tasks. Attention, Perception & Psychophysics, 71(8), 1782–1792. 10.3758/APP.71.8.1782

24. Smith, D. T. (2022). A horizontal–vertical anisotropy in spatial short-term memory. Visual Cognition, 30(4), 245–253. 10.1080/13506285.2022.2042446

25. Muhle-Karbe, P. S., Myers, N. E., & Stokes, M. G. (2021). A Hierarchy of Functional States in Working Memory. The Journal of Neuroscience, 41(20), 4461– 4475. 10.1523/JNEUROSCI.3104-20.2021

26. Cai, Y., Fulvio, J. M., Yu, Q., Sheldon, A. D., & Postle, B. R. (2020). The Role of Location-Context Binding in Nonspatial Visual Working Memory. Eneuro, 7(6), ENEURO.0430-20.2020. 10.1523/ENEURO.0430-20.2020

27. Duncan, D. H., van Moorselaar, D., & Theeuwes, J. (2023). Pinging the brain to reveal the hidden attentional priority map using encephalography. Nature Communications, 14(1), Article 1. 10.1038/s41467-023-40405-8

28. Van Ede, F., Chekroud, S. R., Stokes, M. G., & Nobre, A. C. (2019). Concurrent visual and motor selection during visual working memory guided action. Nature Neuroscience, 22(3), 477–483. 10.1038/s41593-018-0335-6

29. Liu, J., Zhang, H., Yu, T., Ni, D., Ren, L., Yang, Q., Lu, B., Wang, D., Heinen, R., Axmacher, N., & Xue, G. (2020). Stable maintenance of multiple representational formats in human visual short-term memory. Proceedings of the National Academy of Sciences, 117(51), 32329–32339. 10.1073/pnas.2006752117

30. Liu, J., Zhang, H., Yu, T., Ren, L., Ni, D., Yang, Q., Lu, B., Zhang, L., Axmacher, N., & Xue, G. (2021). Transformative neural representations support long-term episodic memory. Science Advances, 7(41), eabg9715. 10.1126/sciadv.abg9715

31. Schneegans, S., & Bays, P. M. (2017). Restoration of fMRI Decodability Does Not Imply Latent Working Memory States. Journal of Cognitive Neuroscience, 29(12), 1977–1994. 10.1162/jocn_a_01180

32. Trübutschek, D., Marti, S., Ueberschär, H., & Dehaene, S. (2019). Probing the limits of activity-silent non-conscious working memory. Proceedings of the National Academy of Sciences, 116(28), 14358–14367. 10.1073/pnas.1820730116

33. Wolff, M. J., Kandemir, G., Stokes, M. G., & Akyürek, E. G. (2020). Unimodal and Bimodal Access to Sensory Working Memories by Auditory and Visual Impulses. The Journal of Neuroscience, 40(3), 671–681. 10.1523/JNEUROSCI.1194-19.2019

34. Gayet, S., Paffen, C. L. E., & Van Der Stigchel, S. (2018). Visual Working Memory Storage Recruits Sensory Processing Areas. Trends in Cognitive Sciences, 22(3), 189–190. 10.1016/j.tics.2017.09.011

35. Harrison, S. A., & Tong, F. (2009). Decoding reveals the contents of visual working memory in early visual areas. Nature, 458(7238), 632–635. 10.1038/nature07832

36. Katus, T., Grubert, A., & Eimer, M. (2015). Electrophysiological Evidence for a Sensory Recruitment Model of Somatosensory Working Memory. Cerebral Cortex, 25(12), 4697–4703. 10.1093/cercor/bhu153

37. Bettencourt, K. C., & Xu, Y. (2016). Decoding the content of visual short-term memory under distraction in occipital and parietal areas. Nature Neuroscience, 19(1), Article 1. 10.1038/nn.4174

38. Zhang, J., Ye, C., Sun, H.-J., Zhou, J., Liang, T., Li, Y., & Liu, Q. (2022). The passive state: A protective mechanism for information in working memory tasks. Journal of Experimental Psychology: Learning, Memory, and Cognition, 48(9), 1235–1248. 10.1037/xlm0001092

39. Rose, N. S., LaRocque, J. J., Riggall, A. C., Gosseries, O., Starrett, M. J., Meyering, E. E., & Postle, B. R. (2016). Reactivation of latent working memories with transcranial magnetic stimulation. Science, 354(6316), 1136–1139. 10.1126/science.aah7011

40. Faul, F., Erdfelder, E., Buchner, A., & Lang, A.-G. (2009). Statistical power analyses using G*Power 3.1: Tests for correlation and regression analyses. Behavior Research Methods, 41, 1149–1160.

41. Delorme, A., & Makeig, S. (2004). EEGLAB: An open source toolbox for analysis of single-trial EEG dynamics including independent component analysis. Journal of Neuroscience Methods, 134(1), 9–21. 10.1016/j.jneumeth.2003.10.009

42. De Maesschalck, R., Jouan-Rimbaud, D., & Massart, D. L. (2000). The Mahalanobis distance. Chemometrics and Intelligent Laboratory Systems, 50(1), 1–18. 10.1016/S0169-7439(99)00047-7

43. Maris, E., & Oostenveld, R. (2007). Nonparametric statistical testing of EEG- and MEG-data. Journal of Neuroscience Methods, 164(1), 177–190. 10.1016/j.jneumeth.2007.03.024

44. Foster, J. J., Sutterer, D. W., Serences, J. T., Vogel, E. K., & Awh, E. (2016). The topography of alpha-band activity tracks the content of spatial working memory. Journal of Neurophysiology, 115(1), 168–177. 10.1152/jn.00860.2015

