## Supplemental information for "Reactivating and Reorganizing Activity-Silent Working Memory Through Two Distinct Mechanisms After Pinging the Brain"

### Supplementary Materials

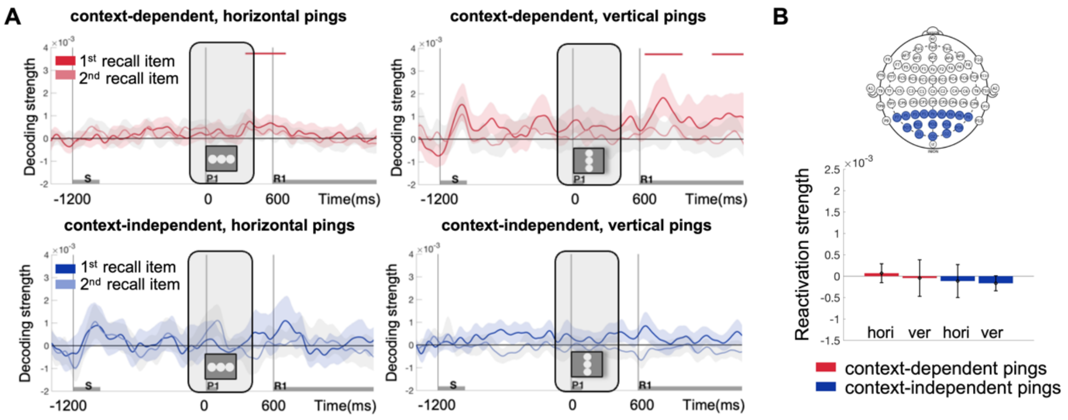


**Supplementary Figure1. Time-resolved decoding and reactivation results based on alpha power activities among posterior areas**. *(A)* Decoding strengths of the first recall item (PMI) under the four ping conditions. The blue and red bars on the top of each plot indicated clusters with significant decoding strengths and the black bars indicated clusters with significant decoding differences between the items ($p < 0.05$). Error shading along the decoding strength line indicated a 95% confidence interval. Gray bars on the X-axis indicated the duration of the sample (S), the first ping (P1), and the first recall (R1). Transparent gray areas with solid borders indicated the time window used to estimate the immediate reactivation strengths (-100 ms ~ 0 ms before the first ping and 100 ms ~ 400 ms after the first ping). *(B)* immediate reactivation strengths of the first recall item (prioritized item, PMI) under the four ping conditions. Error bars indicated the standard error of the mean (SEM).

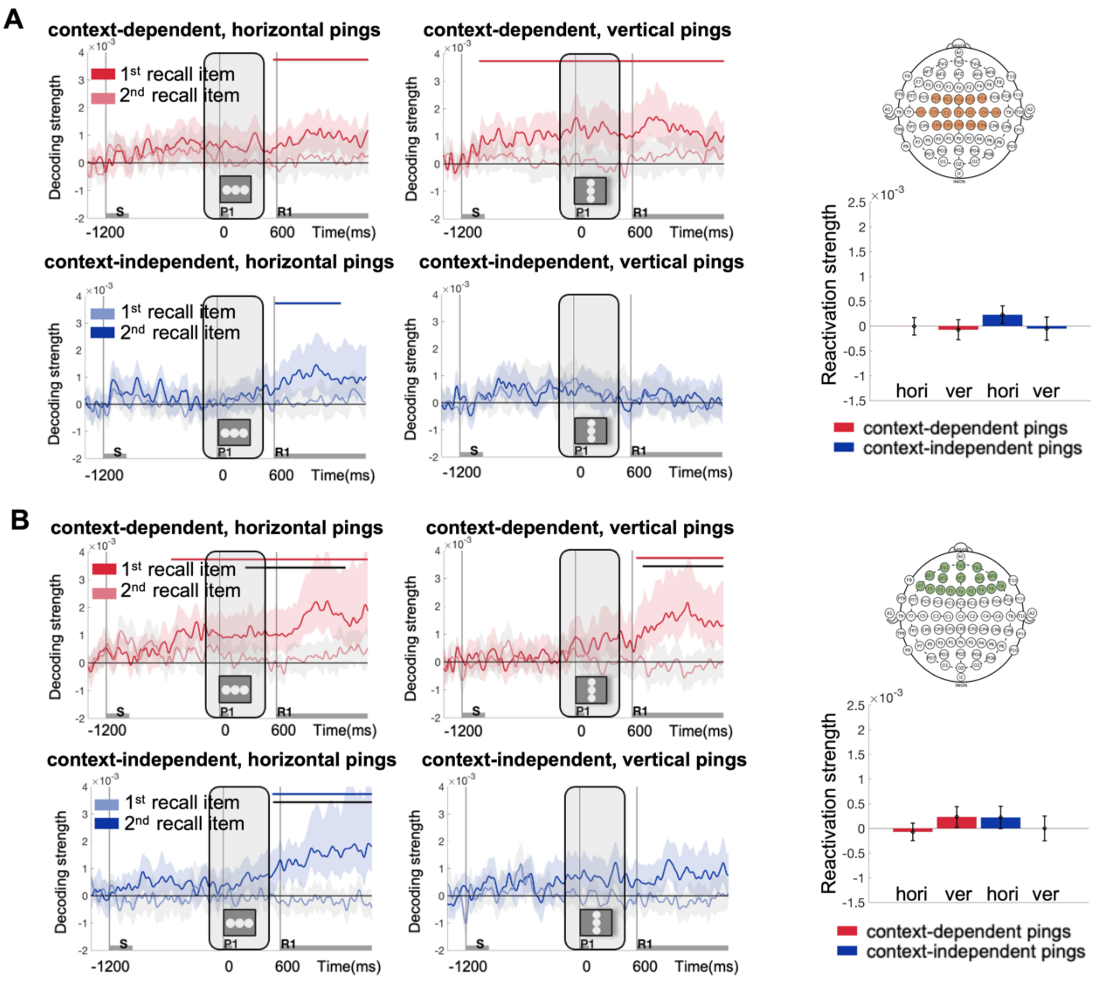


**Supplementary Figure2. Time-resolved decoding and reactivation results based on voltage activities among mid-central (A) and anterior areas (B).** The blue and red bars on the top of each plot indicated clusters with significant decoding strengths and the black bars indicated clusters with significant decoding differences between the two items ($p < 0.05$). Error shading along the decoding strength line indicated a 95% confidence interval. Gray bars on the X-axis indicated the duration of the sample (S), the first ping (P1), and the first recall (R1). Transparent gray areas with solid borders indicated the time window used to estimate the immediate reactivation strengths (-100 ms ~ 0 ms before the first ping and 100 ms ~ 400 ms after the first ping). The lower right plot: immediate reactivation strengths of the first recall item (prioritized item, PMI) under the four ping conditions. Error bars indicated the standard error of the mean (SEM).

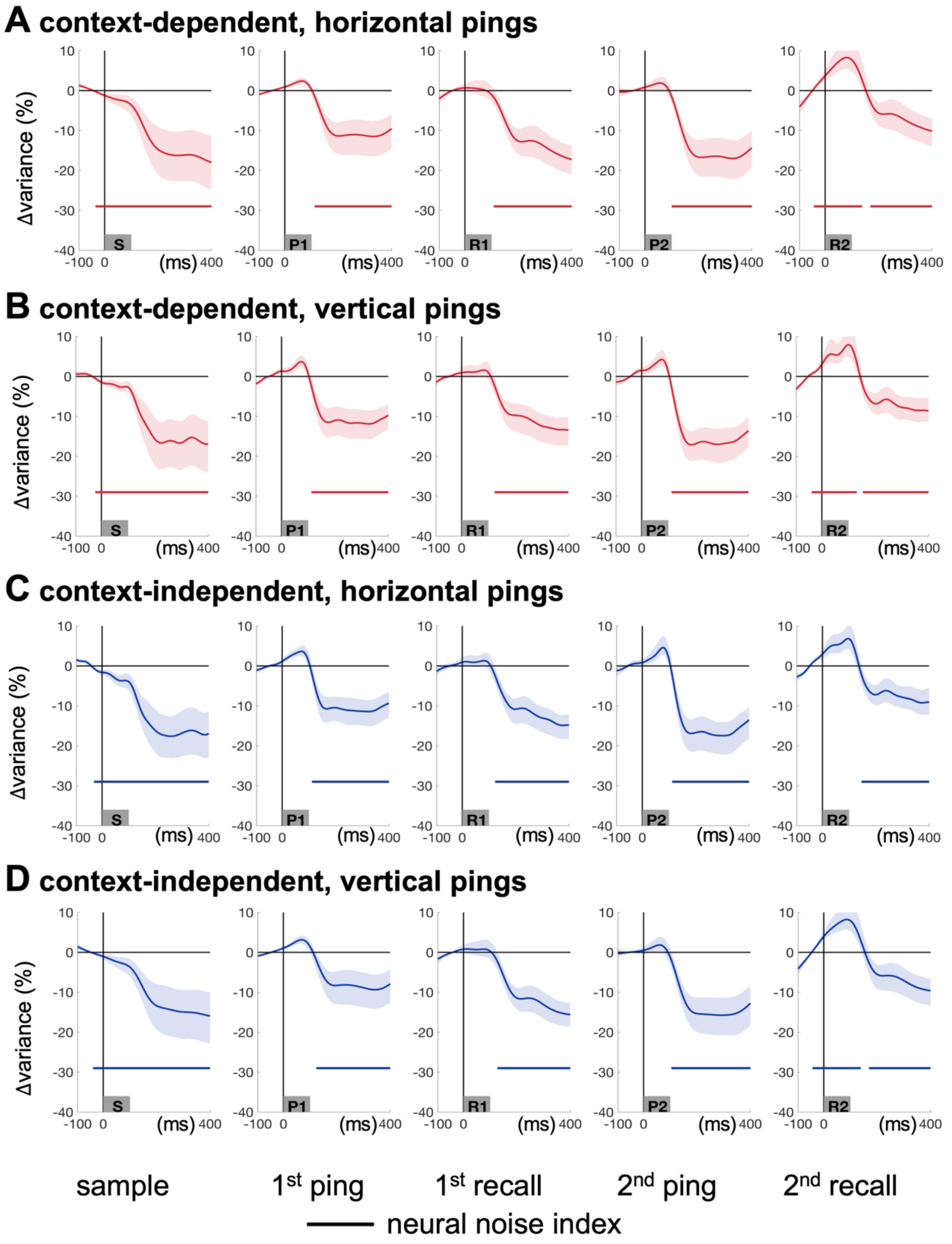


**Supplementary Figure3. Neural noise change after the onset of each visual stimuli in the four ping conditions.** Lines at the bottom of each subplot indicated clusters with significant neural noise reduction compared to the baseline ($p < 0.05$). Error shading indicated a 95% confidence interval of the noise changes. Gray bars on the X-axis indicated the onset of the sample (S), first ping (P1), first recall (R1), second ping (P2), and second recall (R2).
